# Design and application of a knowledge network for automatic prioritization of drug mechanisms

**DOI:** 10.1101/2021.04.15.440028

**Authors:** Michael Mayers, Roger Tu, Dylan Steinecke, Tong Shu Li, Núria Queralt-Rosinach, Andrew I. Su

**Author notes:** equal contribution (Michael Mayers, Roger Tu).

## Abstract

**Motivation:** Drug repositioning is an attractive alternative to de novo drug discovery due to reduced time and costs to bring drugs to market. Computational repositioning methods, particularly non-black-box methods that can account for and predict a drug’s mechanism, may provide great benefit for directing future development. By tuning both data and algorithm to utilize relationships important to drug mechanisms, a computational repositioning algorithm can be trained to both predict and explain mechanistically novel indications.

**Results:** In this work, we examined the 123 curated drug mechanism paths found in the drug mechanism database (DrugMechDB) and after identifying the most important relationships, we integrated 18 data sources to produce a heterogeneous knowledge graph, MechRepoNet, capable of capturing the information in these paths. We applied the Rephetio repurposing algorithm to MechRepoNet using only a subset of relationships known to be mechanistic in nature and found adequate predictive ability on an evaluation set with AUROC value of 0.83. The resulting repurposing model allowed us to prioritize paths in our knowledge graph to produce a predicted treatment mechanism. We found that DrugMechDB paths, when present in the network were rated highly among predicted mechanisms. We then demonstrated MechRepoNet’s ability to use mechanistic insight to identify a drug’s mechanistic target, with a mean reciprocal rank of .525 on a test set of known drug-target interactions. Finally, we walked through a repurposing example of the anti-cancer drug imantinib for use in the treatment of asthma, to demonstrate this method’s utility in providing mechanistic insight into repurposing predictions it provides.

**Availability and implementation:** The Python code to reproduce the entirety of this analysis is available at: https://github.com/SuLab/MechRepoNet

**Supplementary information:** Supplemental information is available at Bioinformatics online.

## Introduction

Drug repositioning, the discovery and development of new indications for previously developed drugs, promises a quicker, less-costly alternative to traditional *de novo* drug discovery (Ashburn and Thor, 2004). Historically, many repositioning candidates have been identified through clinical observations. Computational repositioning leverages statistical modeling to analyze drug-disease combinations quickly in the hopes of streamlining the candidate identification process. Unfortunately, most candidates identified through this method fail to progress beyond *in vitro* studies (Oprea and Overington, 2015). Several approaches for overcoming this shortcoming have specifically addressed increasing predictive performance on a known gold standard through either updated algorithms (Luo *et al*., 2018) or improving quality and quantity of validated examples for training (Brown and Patel, 2018; Shameer *et al*., 2018). Other methods focus on improving interpretability of predictions, aiming to uncover new experimental avenues through testable hypotheses that may improve understanding of a drug’s effect on a disease (Malas *et al*., 2019). Integrating both computational and experimental approaches, so that the results of each inform the other, is considered the best option for progressing the field of drug repositioning (Pushpakom *et al*., 2019).

Methods for computational repurposing models frequently rely on drug-drug and/or disease-disease similarity to produce predictions (Li *et al*., 2016; Li and Lu, 2012; Luo *et al*., 2018). While this information is both important and predictive, a semantic network can describe complex and contextual relationships between a drug and a disease by connecting several individual concepts in a sequence of relationships. These concepts and relationships (collectively a ”path”) can be represented as nodes and edges, respectively, of a knowledge graph *In silico* predictive models built from such knowledge graphs have the potential to inform experimental work through providing interpretable mechanistic explanations. However, these models heavily weight the relatedness of two drugs, through targets or pharmacology, as well as the relatedness of diseases, both carrying significant predictive power (Himmelstein *et al*., 2017; Emig *et al*., 2013; Mayers *et al*., 2019). In cases where there are no known treatments for a disease, or no diseases known to be similar, which is often the case with rare diseases, these patterns lose their utility. Uncovering connections between a drug and a disease that are less immediate may provide additional evidence for drug repositioning. By taking the few mechanistic details that may be available about a rare disease and identifying how a drug might interact with them could provide better insight.

In this work, we created a heterogeneous network with an emphasis on expressing relationships describing a drug’s mechanism of action. This mechanistic repositioning network (MechRepoNet) was then used as the data source to construct a predictive model for drug repositioning using a modified version of the Rephetio algorithm (Himmelstein *et al*., 2017). Edges representing drug-drug or disease-disease similarity were excluded from the model, enriching for paths with mechanistic meaning. The predictions from the resulting model were shown to be highly enriched for both the true mechanisms of action (as expressed in DrugMechDB) and a drug’s annotated protein target. While demonstrating the value of focusing on mechanistic interpretability, this work also underscored a fundamental challenge in computational drug repurposing based on knowledge graphs in that many key mechanistic relationships are not readily available in structured databases.

## Methods

### Integration of data sources to build a mechanistic network MechRepoNet

#### Selection of data sources

We created DrugMechDB, a database that describes drug mechanisms as paths through a heterogeneous knowledge graph (Mayers *et al*., 2020). DrugMechDB is a relatively small database, with just 383 unique concepts (spanning 13 semantic types) and 432 relationships. But these concepts (nodes) and relationships (edges) were assembled into mechanistic paths describing 123 pairs of diseases and drugs. We examined the concept and relationship types contained within the paths of the Drug Mechanism Database (DrugMechDB) to prioritize relationships important to expressing drug mechanisms.

We used DrugMechDB as a roadmap in the creation of MechRepoNet, a large heterogenous network suitable for use as an input knowledge graph for computational drug repurposing. Data sources were evaluated based on their ability to express the semantic relationships found in DrugMechDB in order to construct a heterogeneous network capable of capturing these paths. Wikidata and Reactome were selected as primary data sources due to their high level of breadth, high data quality, and open reuse policies. Nodes and edges from various data sources that follow the schema captured in DrugMechDB and not abundant in Wikidata or Reactome, were prioritized and added into MechRepoNet. In all, eighteen data sources were selected for integration: Wikidata (Vrandečić, 2012), Reactome (Fabregat *et al*., 2018), InterPro (Mitchell *et al*., 2019), Ensembl (Cunningham *et al*., 2019), miRTarBase (Chou *et al*., 2018), ComplexPortal (Meldal *et al*., 2015), Gene Ontology (Ashburner *et al*., 2000), Uber-anatomy Ontology (Mungall *et al*., 2012), Cell Ontology (Diehl *et al*., 2016), Protein Ontology (Natale *et al*., 2017), Disease Ontology (Schriml *et al*., 2019), Human Phenotype Ontology (Köhler *et al*., 2019), NCATS Inxight Drugs (NCATS Inxight: Drugs), DrugBank (Wishart *et al*., 2018), DrugCentral (Ursu *et al*., 2017), Comparative Toxicogenomics Database (CTD) (Davis *et al*., 2019), RheaDB (Morgat *et al*., 2017), and GAUSS (Dutta *et al*., 2019). Although data licensing restrictions prevent the direct redistribution of MechRepoNet, the complete code to assemble the graph is available at https://github.com/SuLab/MechRepoNet/.

#### Data model determination and data integration

To standardize the data in MechRepoNet, concepts in the graph were normalized to the Biolink data model (Mungall *et al*., 2020). To reduce the number of potential features input into the model, relationships (edges) were standardized to a simplified set of predicates. To ensure that edges could contribute to metapaths at multiple levels of granularity, all directed edges were duplicated with the undirected edge equivalent.

Additional data model normalization for nodes was performed to adjust the level of granularity of Wikidata concepts. Wikidata and mygene.info (Xin *et al*., 2016) were used as primary sources to normalize node concepts between resources, augmented by limited string matching for chemical substances. Exact details are provided within the code repository.

### Training a learning model to find mechanistic treatment paths

#### Learning model

We based our analysis on the Rephetio degree weighted path count (DWPC) algorithm (Himmelstein *et al*., 2017), which uses a logistic regression learning model to predict ChemicalSubstance-treats-Disease relationships. Rephetio uses features based on metapaths that join the node types ChemicalSubstances and Diseases in a knowledge graph. “Metapaths” refer to a specific sequence of node types and edge types, whereas “paths” are specific instances of nodes and edges. For example, a three-node and two-edge metapath would be [ChemicalSubstance] - inhibits - [Protein] - causes - [Disease], and a specific instance of this metapath would be the path linking imatinib to the BCR/ABL fusion protein to chronic myelogenous leukemia.

We slightly deviated from the published Rephetio algorithm to ensure computational tractability, since MechRepoNet was significantly larger than the Rephetio knowledge graph and had many more potential metapaths to use as features in the machine learning model. First, whereas Rephetio used both DWPC of each metapath and node degrees of the compound and disease nodes as features in the model, we only used DWPC features. Second, Rephetio used a permutation analysis for feature selection, and we replaced that with a streamlined approach based on feature elimination by weak learners. Network permutation on MechRepoNet would be computationally expensive as it contains over five times the number of nodes (47,031 to 250,035) and four times the number of edges (2,250,197 to 9,652,116). Finally, we removed all metapaths that were primarily based on compound-compound and disease-disease similarity, enriching for metapaths which represent a drug’s mechanism; the remaining metapaths are mechanistic paths. (https://github.com/SuLab/MechRepoNet/blob/main/1_code/13b_Model_Prep_Metapath_Membership_Analysis.ipynb).

#### Training data

A logistic regression was trained on the examples of ChemicalSubstance-treats-Disease triples found within the knowledge graph. There were 69,639 relationships of this type, coming from multiple data sources including Wikidata, NCATS Inxight Drugs, and CTD (Figure 1a). This large number is attributable to the differing chemical and disease vocabulary used by the data sources as well as the use of computational methods for producing network edges including NLP and subclass path contraction.

**Figure 1:**
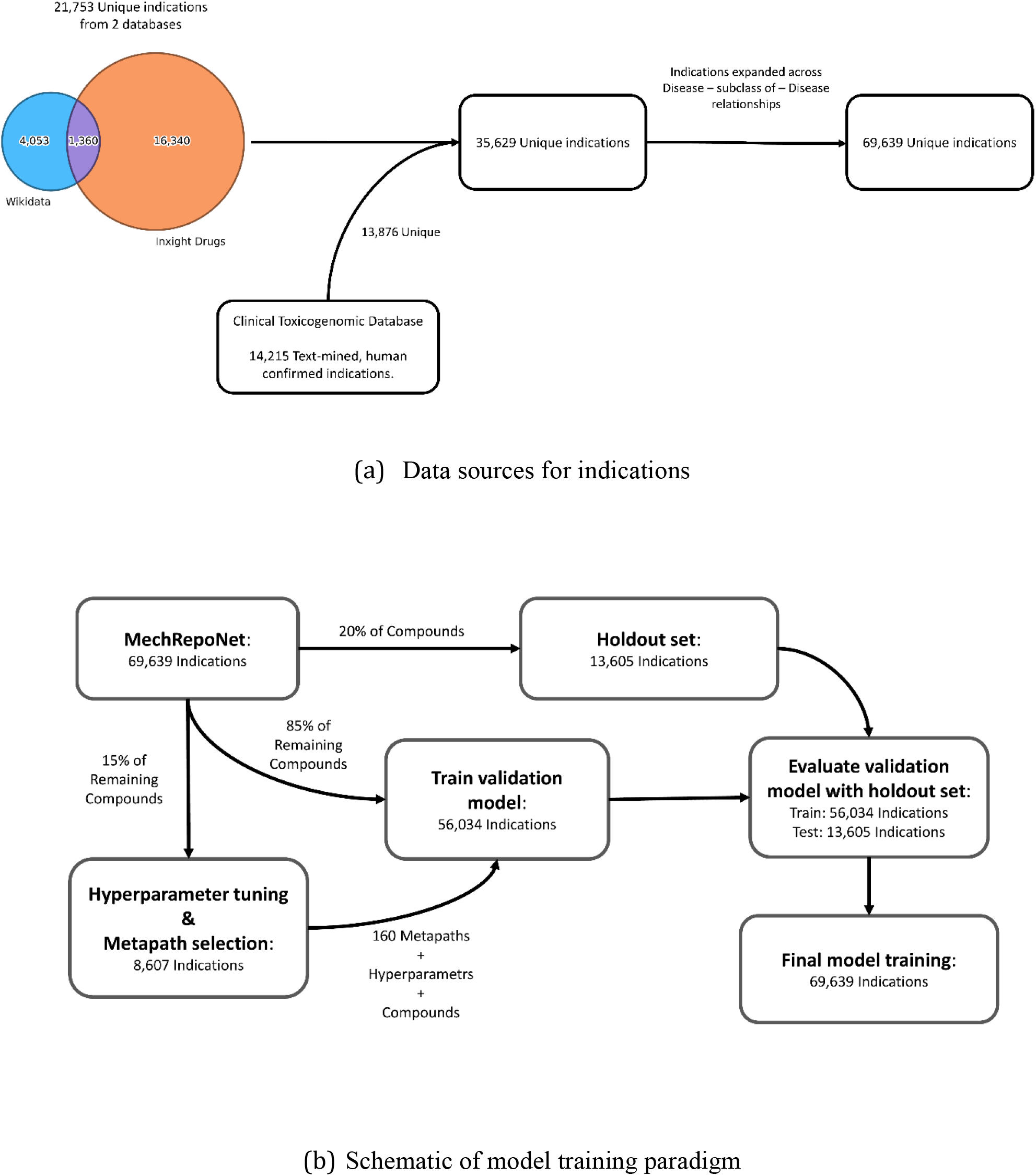
Indication sources and schematic of model training paradigm. a) Indications were initially sourced from two data sources, Wikidata and NCATS Inxight Drugs. Inxight Drugs is a data aggregator that compiles data from several sources including DrugBank and DrugCentral producing many indications. The Clinical Toxicogenomic Database (CTD) contains over 14,000 indications, all originating from text-mining, with human confirmation of the result. Path contraction along the Compound - treats - Disease - inverse subclass of - Disease path produces a total of 69,639 indications. b) Initially, 20% of compounds with known indications were removed and placed in a holdout set. A subset of 15% of the remaining compounds with known indications were used for hyperparameter tuning and metapath selection. The 160 metapaths selected were used to train a model on all indications not within the holdout set for validation of the model. Finally, all indications were used to produce a final model for mechanistic evaluation.

Before any model tuning or training was performed, a subset of 20% of the 11,303 ChemicalSubstances with at least one known indication were selected, and their 13,605 corresponding “treats” edges were removed from the network to be used as a validation set (Figure 1b).

#### Hyperparameter tuning

A subset of 15% of the ChemicalSubstances with known “treats” Disease relationships were selected for hyperparameter tuning, amounting to 1,404 ChemicalSubstances with 8,607 positive training examples (Figure 1b). Negative samples were randomly selected from the set of non-positive ChemicalSubstance-Disease pairs. Negative samples were randomly selected instead of chosen from contraindications to select for no-effect instead of deleterious effects between a ChemicalSubstance and Disease pair. Negative examples were sampled at a rate of 1:1, 10:1, 100:1, and 1000:1 relative to the positive examples to test the model’s sensitivity to the known imbalance between positive and negative treats relationships between drugs and diseases. In all instances of model training, pairs were only utilized if at least one path existed between the compound and the disease. These examples were used for feature selection (see Feature selection below) and selected features were used to tune the DWPC damping exponent (*w*), the elastic net regularization strength (*λ*), and the ratio of *L*_1_ and *L*_2_ penalties (*α*) (Zou and Hastie, 2005). Hyperparameters were tuned using 50 iterations of the Bayesian hyperparameter optimization method (Bergstra *et al*., 2013). Optimization was performed to maximize a linear combination of the area under the receiver operator characteristic and precision recall curves, while minimizing the variance in these values across folds.

#### Feature selection

The fifty-five unique metapaths that were found in DrugMechDB were all included as features in our model. In addition, a voting strategy was employed between six different feature selection methods, each tasked with selecting 500 metapaths. Each feature selection method obeyed one of the following rules: Features with the largest magnitude of correlation coefficient with respect to the target, top features after a *χ*^2^ test between features and targets, a recursive feature elimination of the smallest 10% of features in a weak regression classifier with *L*_2_ regularization, the features with largest magnitude of coefficient from a weak regression classifier with *L*_1_ regularization, the largest feature importance values from a weak random forest classifier, and the largest feature importance values from a weak gradient boosted decision tree. Each selected feature from the 6 methods were considered a vote and features with 4 or more votes were kept for the training model. Code describing this full feature selection process is available on github (https://github.com/SuLab/MechRepoNet/blob/main/1_code/13c_Model_Prep_Hyperparam_tuning.ipynb). This process resulted in fewer than 200 of the 8,284 mechanistic metapaths being selected as features for training. Of the 55 metapaths that were included in our model based on DrugMechDB paths, 21 were selected as features by elastic net regularization, and 12 had positive coefficients for the final MechRepoNet model.

#### Performance evaluation

A model was trained on all the “treats” relationships in the network, less the holdout set as described in the training data section (Figure 1b). This holdout set, along with a sampling of non-positive relationships, was used to validate the learning model built from the data within the network.

## Results

### Analyzing DrugMechDB for relevant relationships

Our first task was to build a heterogeneous network capable of expressing drug mechanisms. To help direct the creation of our network, we created DrugMechDB, a manually curated database of drug mechanisms. DrugMechDB consisted of 123 mechanistic paths for 109 unique indications between 106 drugs and 86 diseases. This dataset spanned a diverse cross-section of drug classes and disease areas (Supplemental Figure 1). The most common edge type in DrugMechDB joins Drug entities with Protein entities, with 94 of the 123 paths containing a relationship of this form (Table 1). Other common edge types join Biological Processes with Disease, and Proteins with Biological Processes.

**Table 1:**
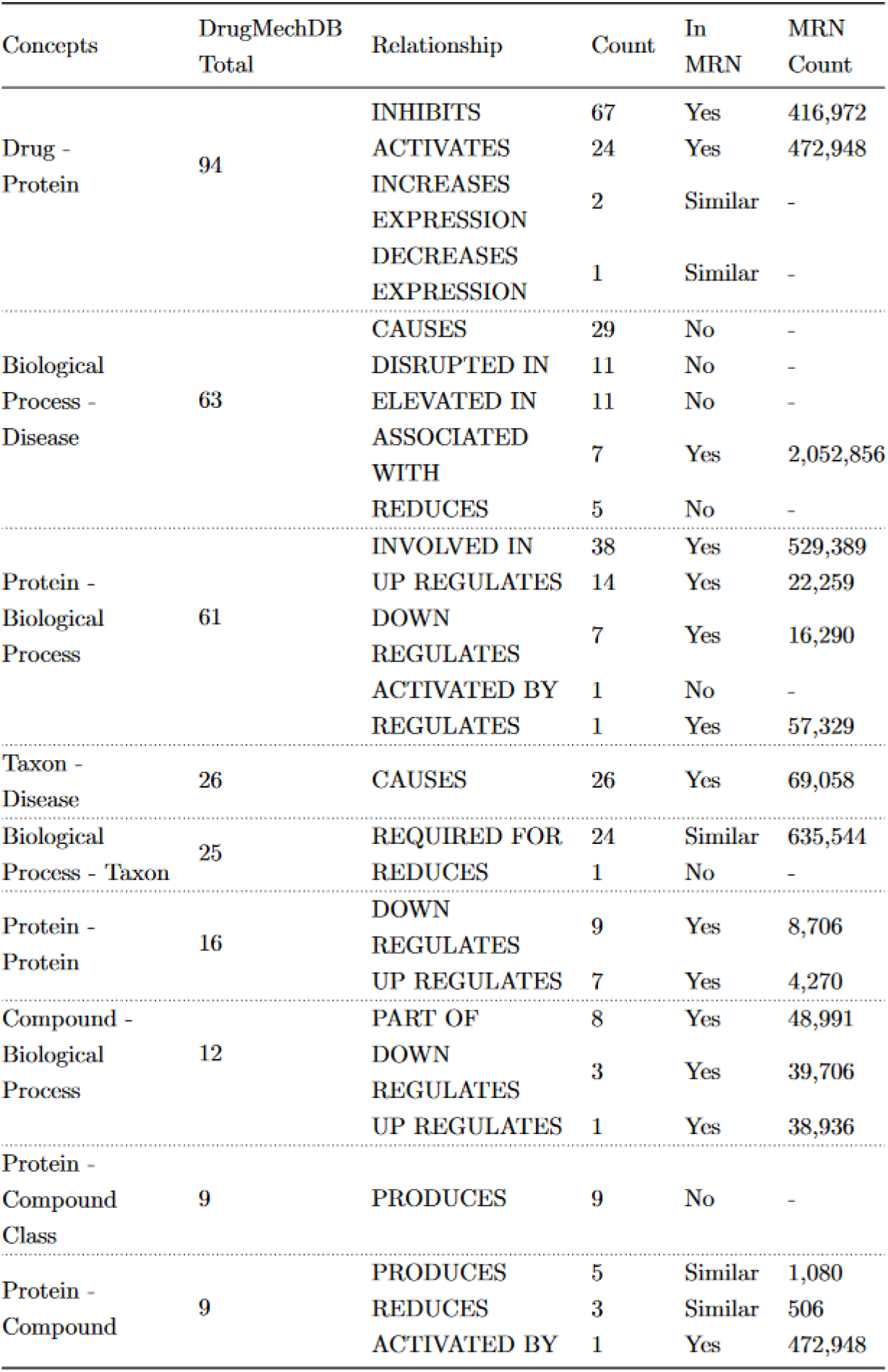
Counts of top relationship types in DrugMechDB. The most common pairings of concepts found in DrugMechDB and the different relationships that connect them. Whether or not these concepts are expressed in MechReopNet is noted, with “Similar” meaning either different semantics are used with similar meaning (e.g. “Down Regulates” instead of “Inhibits”), or that relationship exists, but something about it is fundamentally different (e.g. Protein Regulates Protein relationships without specific up or down directionality). MRN = MechRepoNet.

### Building a network to express drug mechanisms: MechRepoNet

Based on the analysis of mechanistic paths in DrugMechDB, we integrated 18 publicly available data sources to produce a new mechanistic repositioning heterogeneous network (MechRepoNet) (Figure 2a). MechRepoNet contains 250,035 concepts of 9 different semantic types, and 9,652,116 unique edges of 68 semantic types (where the semantic type of an edge is defined by a subject node type, an object node type, and a predicate) (Figure 2b). Most concepts in MechRepoNet are classified as MacromolecularMachine, which includes Genes, Proteins, RNA Products, and Complexes (Table 2). The most common edge type connects a BiologicalProcessOrActivity to a Disease through a general association, with 2,052,856 examples, representing 21.3% of the edges in MechRepoNet (Table 1).

**Figure 2:**
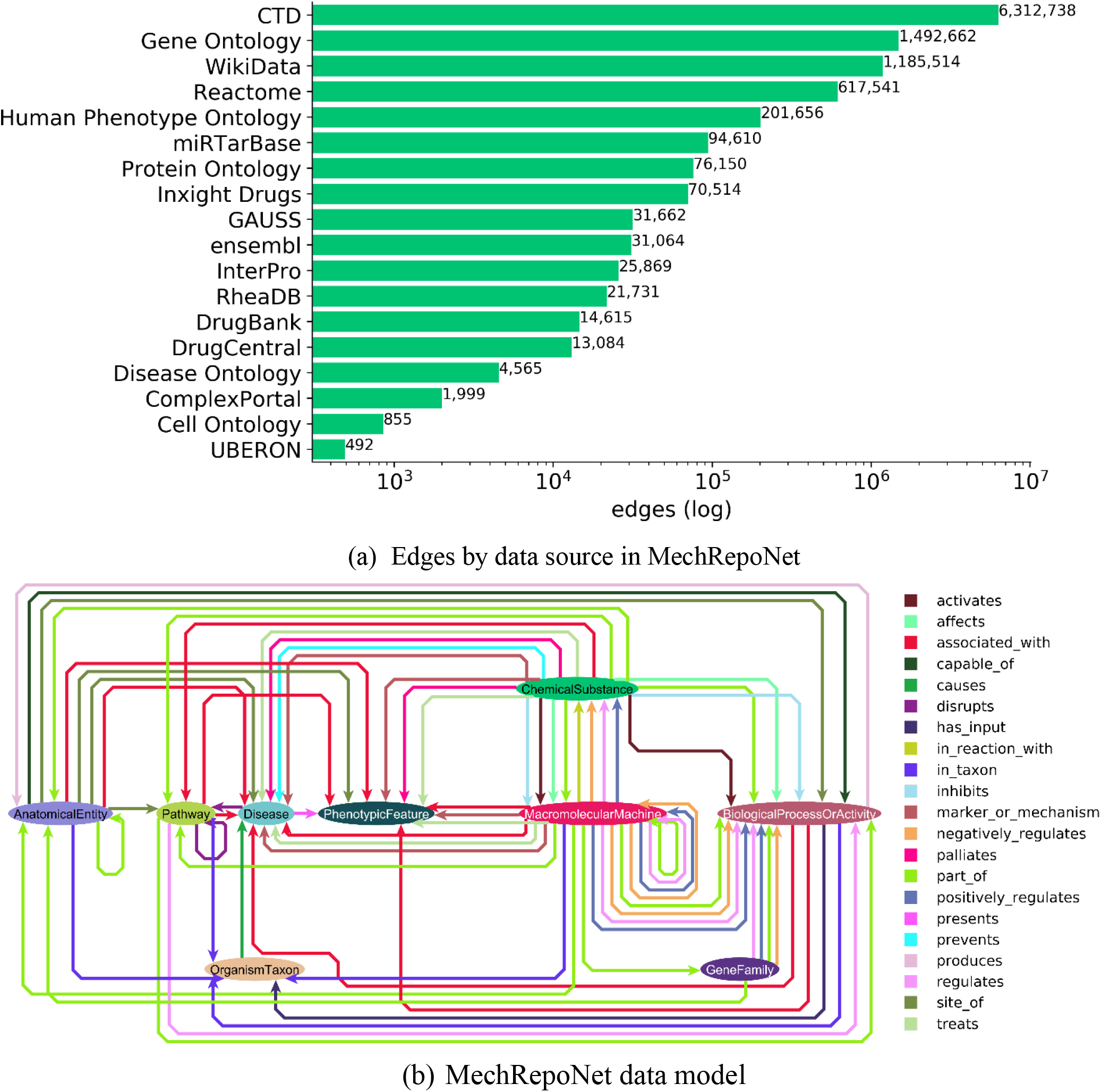
Data within MechRepoNet. (a) The number of edges provided by each data source integrated into MechRepoNet. (b) A graphic representation of the MechRepoNet data model.

**Table 2:** Concept types in MechRepoNet.

| Node type | Abbreviation | Identifier Sources | Count |
| --- | --- | --- | --- |
| AnatomicalEntity | A | GO, UBERON, CL | 3,580 |
| BiologicalProcessOrActivity | BP | GO | 21,065 |
| ChemicalSubstance | C | CHEBI, IKEY, MESH, UNII, PCID, ChEMBL, WD, DB | 40,766 |
| Disease | D | DOID, MESH, MONDO, OMIM, WD, UMLS | 13,961 |
| GeneFamily | GF | InterPro | 10,807 |
| MacromolecularMachine | G | NCBIGene, WD, UniProt, RNAC, REACT, CPX, ENSG, PR | 136,991 |
| OrganismTaxon | T | NCBITaxon, WD | 12,074 |
| Pathway | PW | REACT, WP, KEGG, WD | 5,363 |
| PhenotypicFeature | P | HP, WD, MESH, OMIM | 5,428 |

While MechRepoNet contains many of the concept types found in DrugMechDB, there are a few differences (Table 2). Most notably, we normalized concept types to the Biolink data model, a standardized hierarchy of biomedical entity classes, to increase interoperability between data sources (Mungall *et al*., 2020). Relationships in MechRepoNet were mapped to a flat list of interaction types (Table 3). In general, the relationships in DrugMechDB are more specific than those in MechRepoNet, primarily because data sources use imprecise semantics when describing relationships.

**Table 3:** Relationship types in MechRepoNet.

| Relationship type | Abbreviation | Count | Relationship type (cont.) | Abbreviation (cont.) | Count (cont.) |
| --- | --- | --- | --- | --- | --- |
| activates | a | 511,884 | negatively_regulates | nr | 21,439 |
| affects | af | 986,843 | palliates | pl | 2,519 |
| associated_with | aw | 4,488,453 | part_of | po | 1,252,343 |
| capable_of | co | 513 | positively_regulates | pr | 32,319 |
| causes | co | 69,058 | presents | ps | 66,965 |
| disrupts | d | 541 | prevents | pv | 1,318 |
| has_input | hi | 10,905 | produces | p | 1,008 |
| in_reaction_with | rx | 152,741 | regulates | rx | 215,855 |
| in_taxon | it | 1,208,845 | site_of | so | 31,791 |
| inhibits | in | 456,678 | treats | t | 75,798 |
| marker_or_mechanism | m | 64,300 |  |  |  |

We next examined the degree to which DrugMechDB paths could be found within MechRepoNet. We found that 92.2% of the 383 unique concepts in DrugMechDB are also contained within MechRepoNet (Figure 3a). Most of the missing concepts are either proteins from infectious taxa, or the taxa themselves (Supplemental Table S1). Evaluating relationships as simple edges (ignoring edge predicates), we found that 167, or 45% of the 369 unique node parings found in DrugMechDB are also contained with MechRepoNet (additional details in Figure 3b). Of the 202 missing pairings, 58 are due to one or both concepts not being present, whereas the other 144 have both concepts present, but the connection between them is absent. Finally, we found that 20 mechanistic paths from DrugMechDB are fully expressed in MechRepoNet, and another 27 only have one edge missing (Figure 3c).

**Figure 3:**
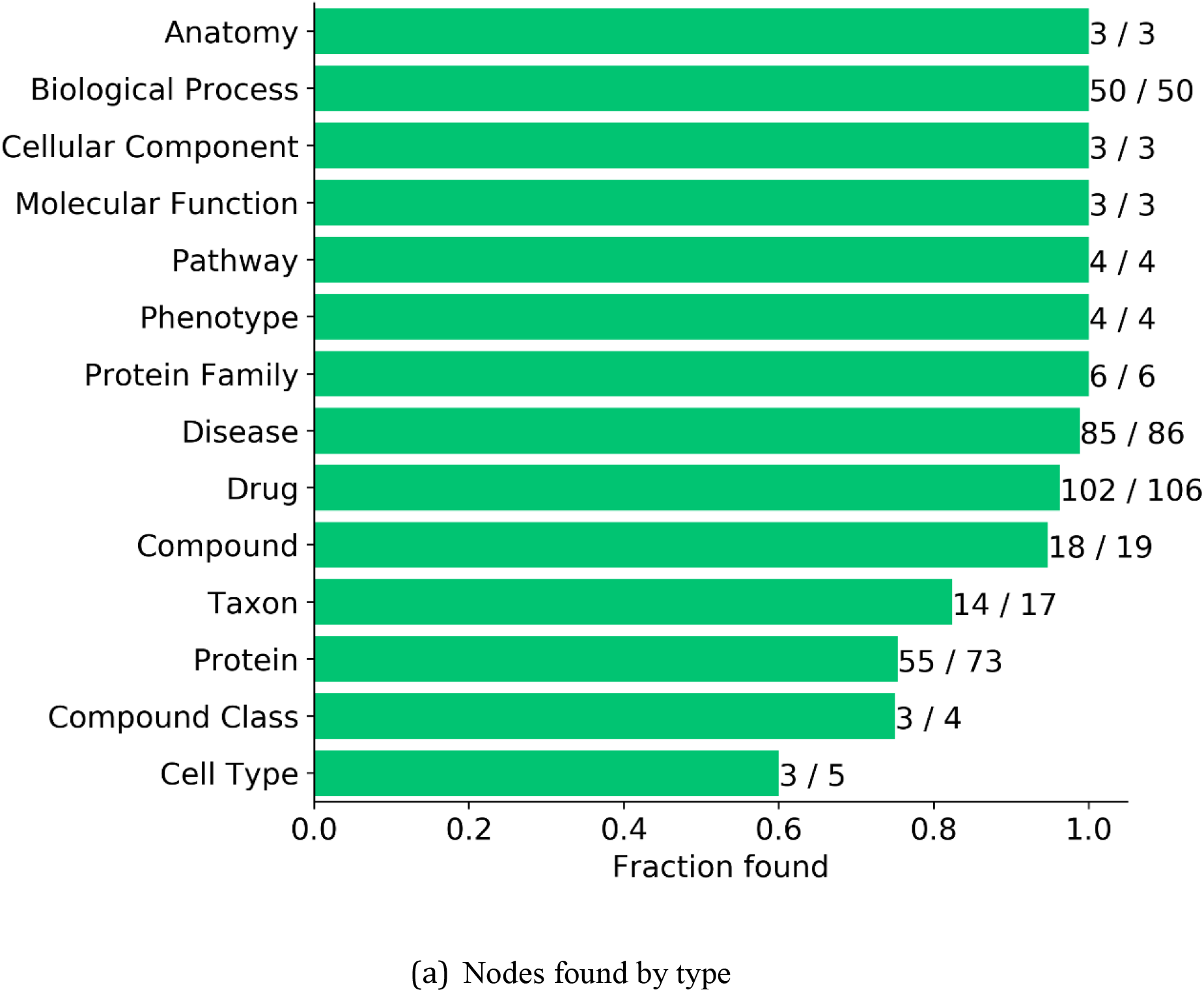
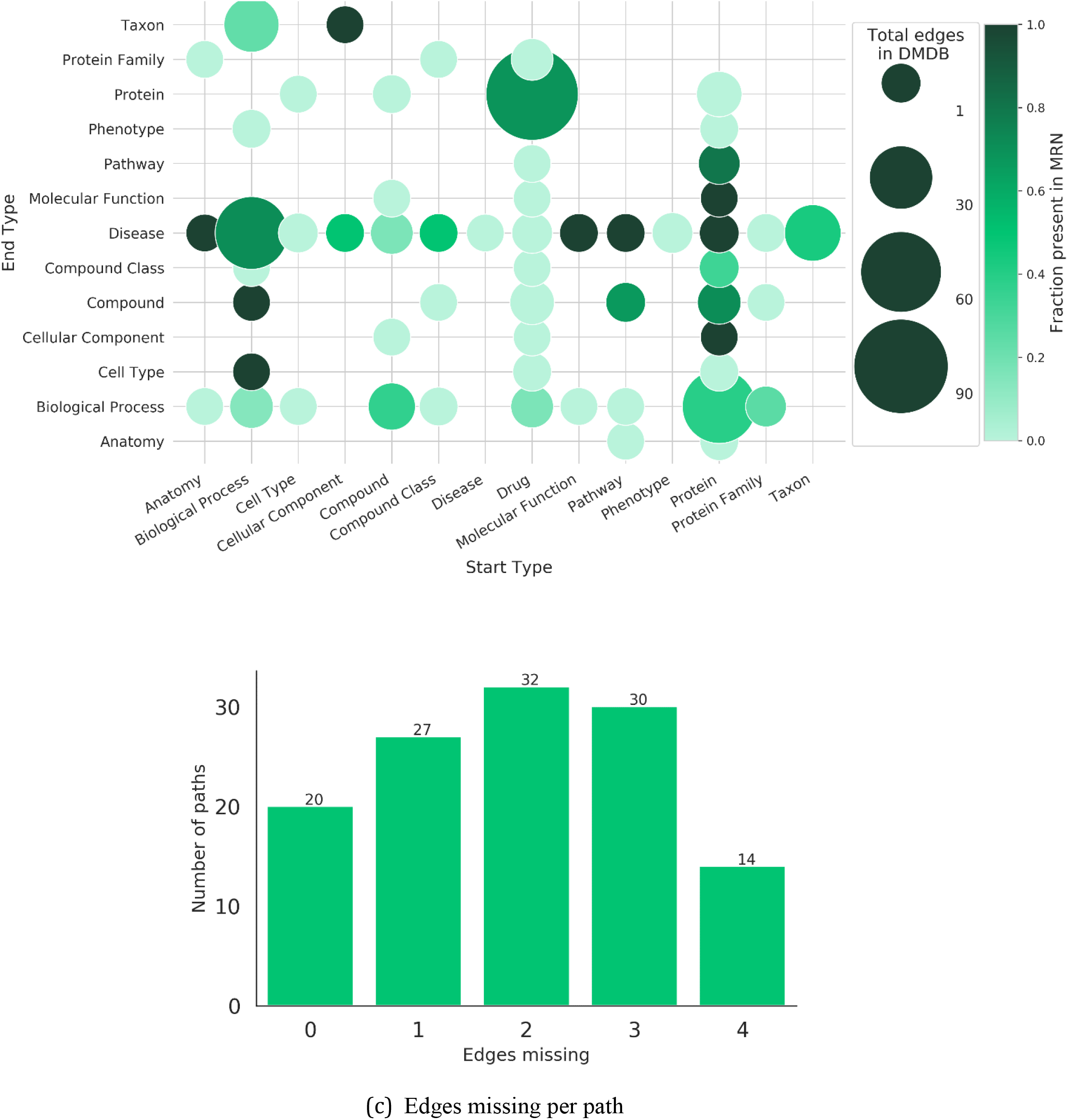
Comparison of concepts, edges and paths found in DrugMechDB and MechRepoNet. (a) Number of unique concepts in DrugMechDB, represented in MechRepoNet by type. (b) Analysis of edges in DrugMechDB and comparison to MechRepoNet. Bubble size is proportional to the number of edges in DrugMechDB, and darker bubbles indicate greater percentage overlap between DrugMechDB and MechRepoNet. (c) Number of paths with 0 to 4 edges missing from MechRepoNet for each of the 123 DrugMechDB paths.

### Metapaths for feature extraction

MechRepoNet contains 17,277 unique metapaths of length 2, 3, and 4 between a ChemicalSubstance and a Disease. Removing those that express compound-compound or disease-disease similarity (see Methods) resulted in 8,284 remaining metapaths. Even though these metapaths may exist within the data model of a network, it is not guaranteed that there are paths within the network that follow that metapath structure. Of the 8,284 mechanistic metapaths, 7,012 (84.6%), were found to be present in MechRepoNet. A feature selection voting scheme (see Methods) filtered this list down to 105 metapaths. After adding the 55 metapaths found in DrugMechDB, we ended up with a final set of 160 metapaths to use as features for training our learning model.

### Learning model results

Here, we characterize the performance of our mechanistically focused analysis. Regardless of negative to positive sampling ratio, the models performed well on the validation set (see Figure 1 for experimental setup) with the area under the curve for the receiver operating characteristic (ROC AUC) of 0.83 (for the 100 to 1 negative to positive ratio) which indicates a high rate of true positives for each false positive result (Figure 4a). Since ROC AUC results can be skewed by the class imbalance from our sampling scheme (100 to 1 negative to positive ChemicalSubstances to Disease pairs), we also computed the AUC for the precision-recall curve (PRC AUC). The PRC AUC was much lower at 0.03, which is expected in a complicated task with imbalanced classes (Figure 4b).

**Figure 4:**
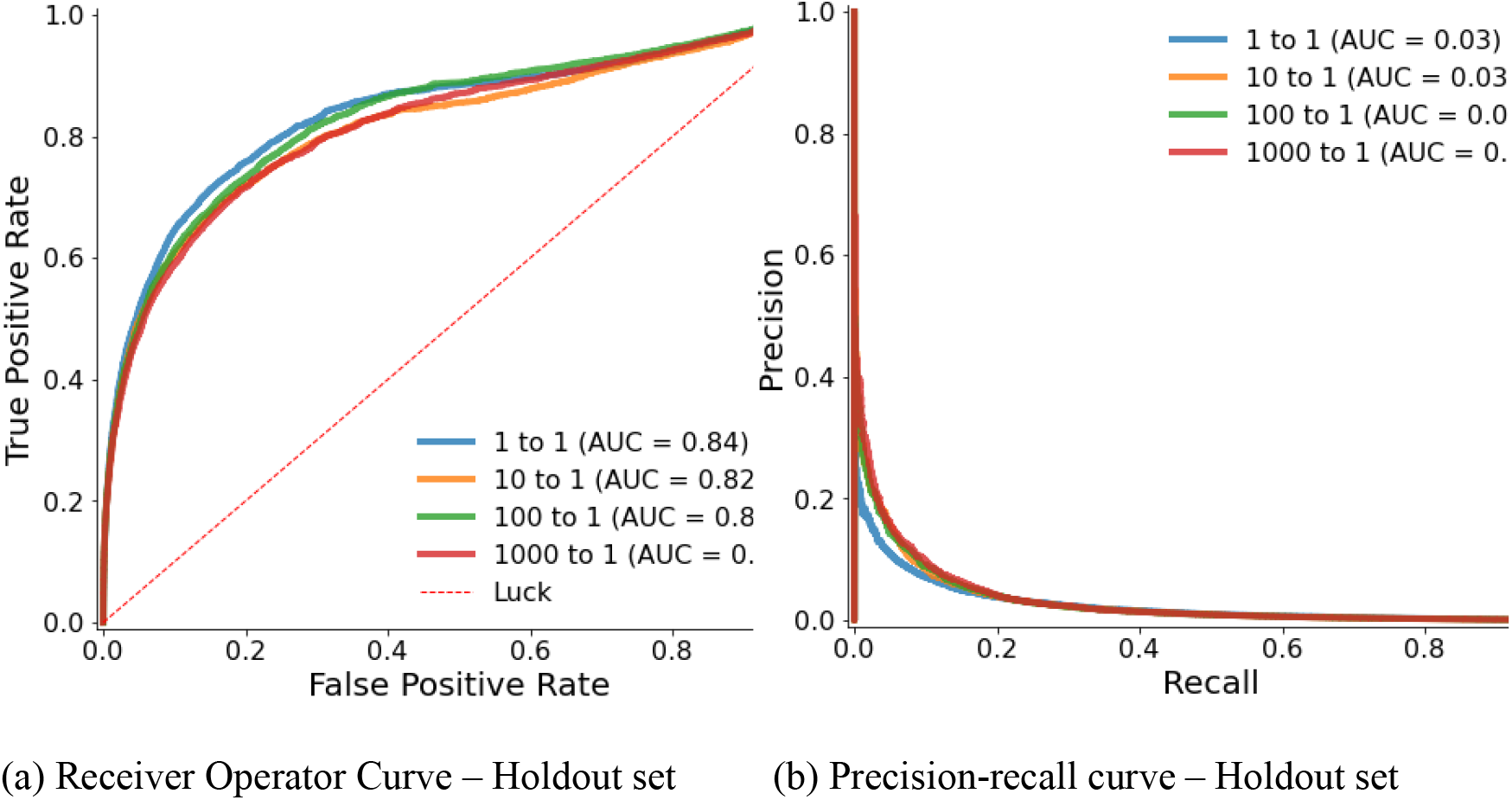
MechRepoNet model validation performance. Model’s performance on holdout set of 20% of indications (a) ROC & (b) precision recall for 1:1, 10:1, 100:1 and 1000:1 negative to positive ChemicalSubstance to Disease pairs. The green line is the chosen model and the red y=x denotes a coin flip.

The ROC AUC and the corresponding AUC for the precision-recall curve (PRC AUC) did not change significantly when sampling ratio increases or decreases from the initial 100 to 1 negative to positive sampling scheme. The minor effect on ROC AUC and PRC was attributed to the hyperparameter tuning step, which adjusts the model hyperparameters to each models sampling scheme, causing the six weak learners to choose slightly different features to push to elastic net regularization.

When varying the ratio of negative to positive sampling, the distribution of prediction probabilities shifted lower as the negative to positive ratio increased (Supplemental Figure 2). However, the ranked performance of the individual models did not change the prediction hits@k significantly (Supplemental Table S2). This further provides evidence that while the increased negative to positive sampling ratio may affect the prediction probability of the model generated, the models maintained their performance.

While these performance metrics indicate that there is still plenty of room for algorithmic improvements (and superior methods may even already exist), these results are sufficient to examine the interpretability of predictions proposed by this metapath-based model. Of the 160 features used to train the model, 89 were assigned positive coefficient values by the regression model, making their associated metapaths meaningful in the mechanistic descriptions of the prediction. From the 55 DrugMechDB metapaths, 27 were selected by the model with positive coefficients (Table 4). The selected features contained all the graph’s 9 node types, with MacromolecularMachine nodes appearing most frequently after ChemicalSubstance and Disease nodes which are guaranteed to be present in every metapath (Figure 5).

**Table 4:**
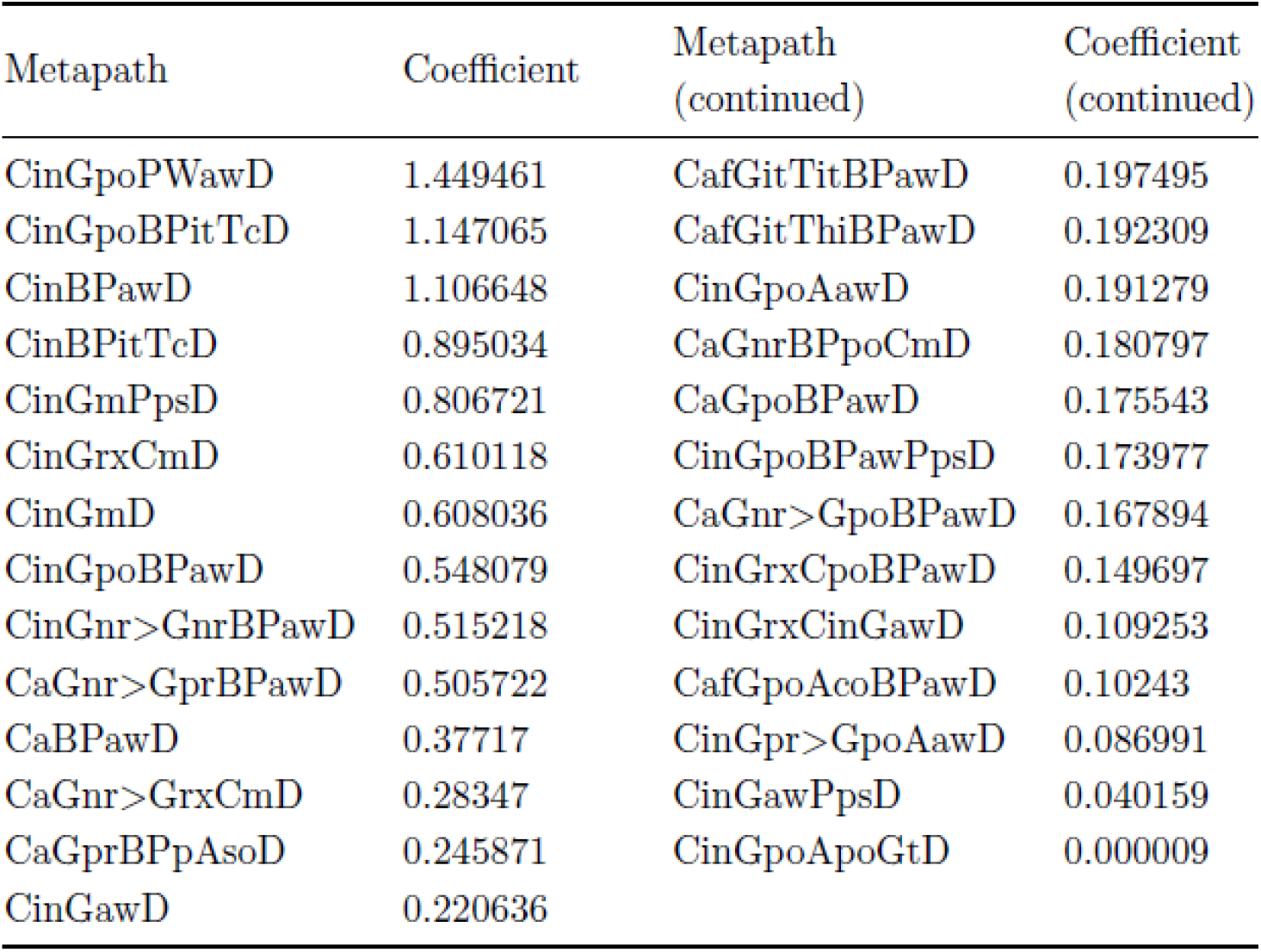
Positive model coefficients for DrugMechDB metapaths. This table contains all the metapaths that were assigned positive coefficients by the logistic regression as well as the coefficient value. Node abbreviations are in capital letters, and relationship abbreviations are in lower case. Node abbreviations are capitalized while edge abbreviations are lower case. See Tables 2 and 3 for node and edge abbreviations. For example, the ‘CinGpoPWawD’ indicates a metapath corresponding to ChemicalSubstance -inhibits-MacromolecularMachine -part of-Pathway -associated with-Disease.

**Figure 5:**
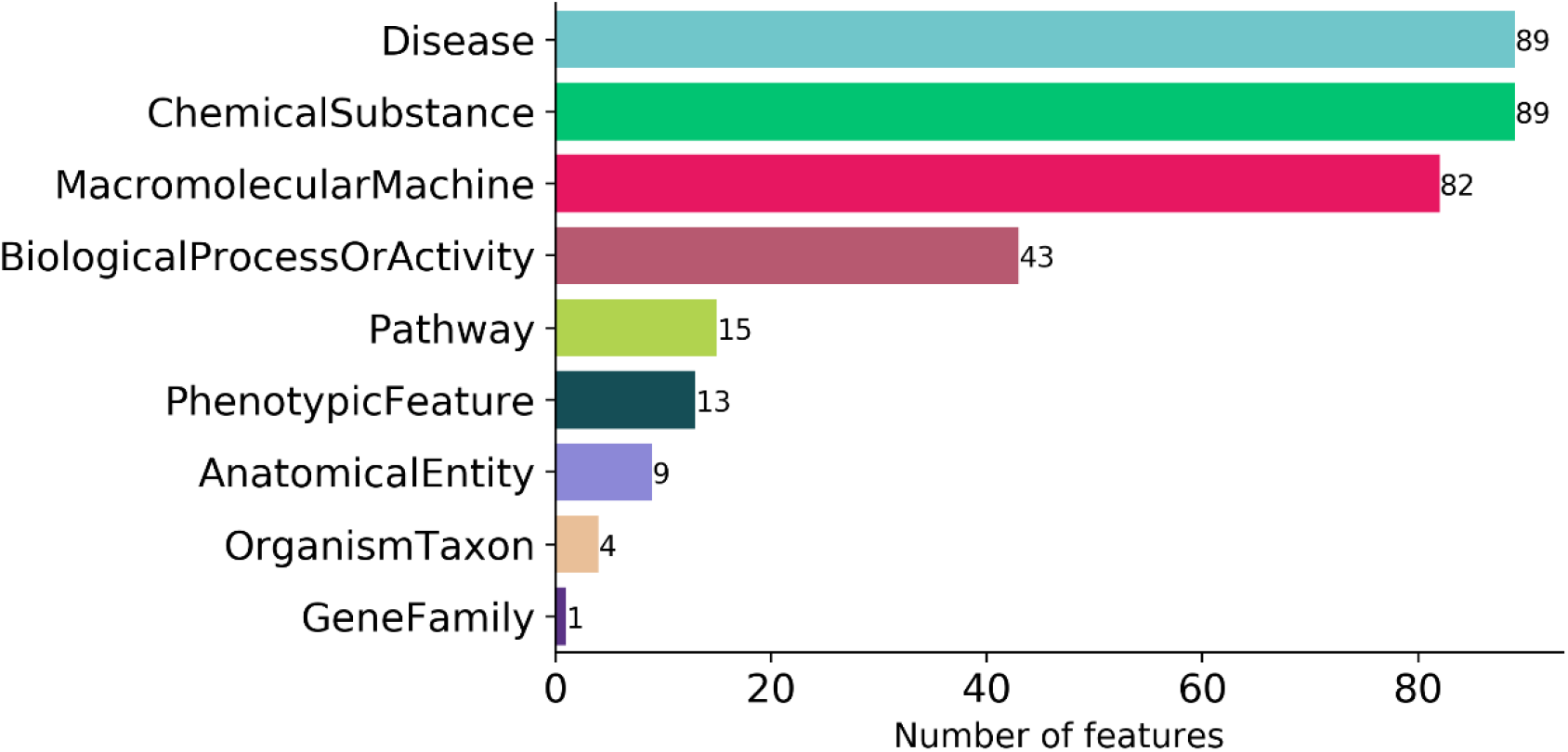
Feature selected concepts. Number of selected features passing through each concept type.

DrugMechDB paths found in MechRepoNet were utilized to evaluate the model’s selected metapaths. Of the 20 DrugMechDB paths, the model identified 12 instances where the corresponding MechRepoNet metapath had a positive coefficient. The coefficient values of the 12 metapaths were compared against other metapaths selected by the model (that do not exist in DrugMechDB). Metapaths with higher coefficients are better predictors than those with lower coefficients. Of the 12 MechRepoNet metapaths, 11 had predicted metapaths above the 90^th^ percentile of all paths (Table 5). In some cases, these rankings may be high enough to quickly support a repurposing prediction, but in others there are still hundreds, or thousands of paths ranked above the mechanistic path in DrugMechDB, making the mechanistic path unlikely to be discovered. These cases present opportunities for further curation efforts to improve metapath predictive performance. For this reason, several filtering strategies can be employed to enrich for these paths.

**Table 5:** DrugMechDB paths as ranked by the repurposing model. DrugMechDB paths found in MechRepoNet and their rankings among all paths between the drug disease pair of an indication. Rankings of subset paths through the drug’s known mechanistic target are also examined (DMDB stands for DrugMechDB).

| Drug | Disease | MRN<br>Paths<br>between<br>Drug &<br>Disease | MRN<br>Predicted<br>Path Rank | Path<br>Percentile | Drug's<br>Target | Paths<br>through<br>Target | MRN<br>Predicted<br>Target<br>Path Rank | Target<br>Path<br>Percentile | Metapath |
| --- | --- | --- | --- | --- | --- | --- | --- | --- | --- |
| imatinib<br>methane<br>-sulfonate | chronic<br>myeloid<br>leukemia | 1,236,778 | 23 | 99.99% | ABL1 | 1,165 | 1 | 99.91% | CafGmD |
| terguride | pulmonary<br>hypertension | 16,523 | 1 | 99.99% | HTR2B | 1,104 | 1 | 99.91% | CafGmD |
| glutethimide | insomnia | 88,253 | 197 | 99.78% | GABRA1 | 143 | 10 | 93.01% | CafGpoBPawD |
| mosapramine | schizophrenia | 4,410 | 15 | 99.66% | DRD2 | 1,792 | 5 | 99.72% | CinGpoPWawD |
| etizolam | anxiety<br>disorder | 42,582 | 210 | 99.51% | GABRA1 | 280 | 28 | 90.00% | CafGpoBPawD |
| vinpocetine | dementia | 291,650 | 2,612 | 99.10% | PDE1A | 134 | 21 | 84.33% | CinGpoAawD |
| (R)<br>-duloxetine | major<br>depressive<br>disorder | 36,692 | 458 | 98.75% | SLC6A4 | 243 | 39 | 83.95% | CafGpoBPawD |
| landiolol | atrial<br>fibrillation | 1,218 | 25 | 97.95% | ADRB1 | 1,005 | 21 | 97.91% | CafGpoBPawD |
| metolazone | hypertension | 13,044 | 296 | 97.73% | SLC12A3 | 646 | 74 | 88.54% | CinGpoBPawD |
| (R,R)<br>-tramadol | pain | 172,613 | 6,822 | 96.05% | OPRM1 | 951 | 290 | 69.51% | CafGpoBPawD |
| gepirone | major<br>depressive<br>disorder | 603 | 34 | 94.36% | HTR1A | 304 | 24 | 92.11% | CafGpoBPawD |
| trimethadione | absence<br>epilepsy | 409 | 244 | 40.34% | CACNA1G | 141 | 83 | 41.13% | CafGpoBPawD |

The most basic filtering strategy is to filter by known drug targets. Most drugs have only a small number of known targets, so filtering this way is straightforward. When filtering these paths by the known mechanistic targets, 9 of the 12 examples are in the top 50 paths, with 4 of 12 in the top 10 paths, making them easy for an expert to identify from the total list of paths (Table 5). The high rankings of the mechanistic paths that pass through known drug targets means that a drugs mechanism for an unknown disease could potentially be found if known targets are used as a filtration step. However, when a true mechanistic target for a drug is not known the mechanistic targets identified from the resulting paths could also be used to identify candidate targets.

### Concept weighting to identify drug targets

The weightings used to rank paths from a drug to a disease can also be used to determine the importance of any single concept connecting them by summing the weights of all the paths containing a given concept. This method could have utility in prioritizing the target of a drug in the treatment of a disease. To evaluate this method, a set 1,124 of known drug-target-disease triples were curated from DrugCentral and compared to either all concepts or only the subset of gene targets, in the paths that connect the drug-disease pair. Examining the percentile rank of known mechanistic targets against all other concepts, most of the known targets ranked in the 90th percentile (Figure 6a). When limiting the comparison to only other gene targets, the percentile rank for the known target was less favorable, producing a distribution that approaches uniform (Figure 6a). However, as some drugs only have a very limited number of potential targets, the absolute ranking of the known target may be informative. Examining the absolute rank of mechanistic targets, when compared to all potential target genes, found 944 of the 1,124 in the top 100, and 637 of those in the top 10 (Figure 6b). The mean reciprocal rank of the known targets, when taking only the highest ranked target for a given drug-disease pair, was found to be 0.525.

**Figure 6:**
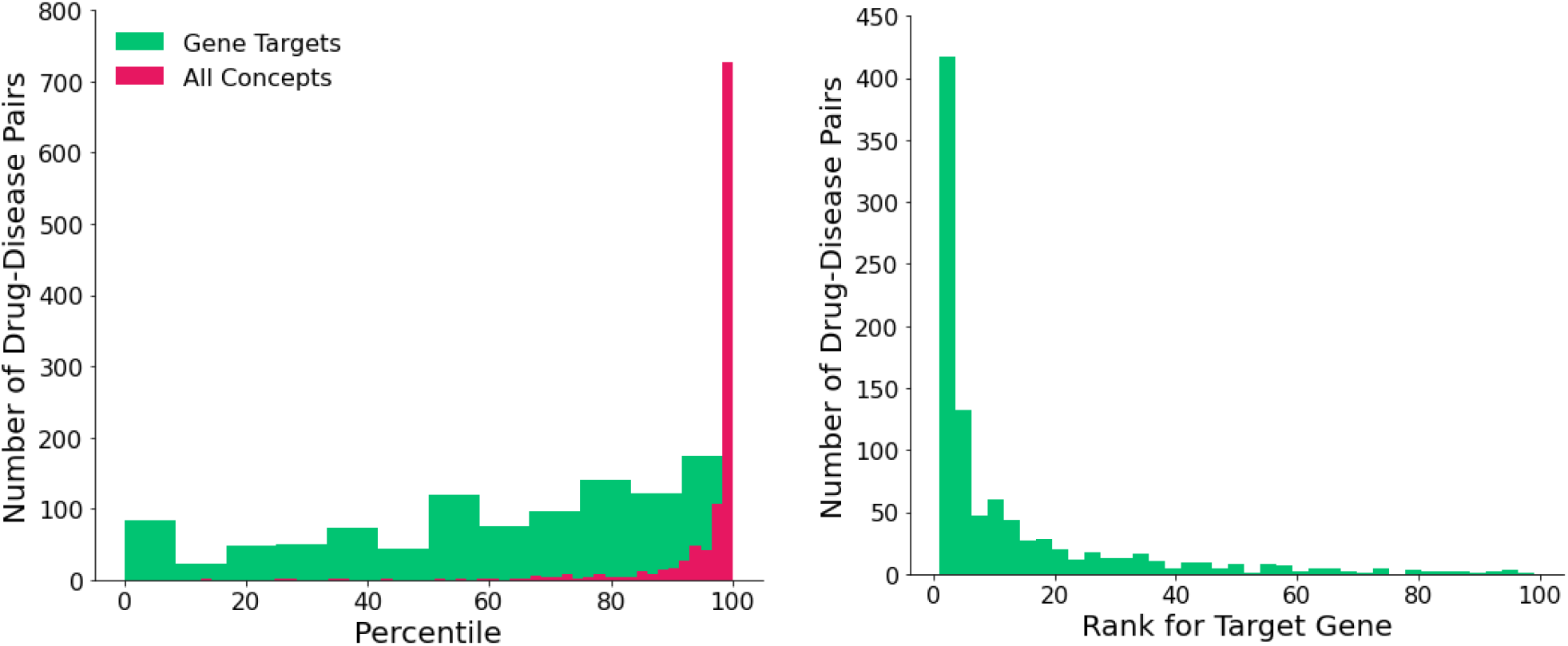
Known target ranking results. (a) Percentile rank of known mechanistic targets of a drug as compared to either all concepts connecting a drug and disease, or only potential gene targets connecting the two. (b) Absolute rank of known mechanistic targets of a drug when compared to other potential gene targets of the drug.

### Case Study: Explaining the link from imatinib to asthma

In 2017 Cahill *et al*. reported that imatinib, a tyrosine kinase inhibitor used to treat chronic myelogenous leukemia, could potentially be repositioned to treat asthma, with promising preliminary results (Cahill *et al*., 2017). We wanted to determine whether the MechRepoNet model was capable of both predicting and explaining this potential repositioning result. The probability score returned by the model for imatinib to asthma was 0.143. However, the absolute probability score in this sense does not have a lot of meaning, and ranking is much more important. The ranking of the imatinib to asthma indication, compared the over 90 million potential drug-disease combinations, was in the top 250,000 putting it in the 99.7th percentile. In terms of the drug and disease specific percentiles, imatinib was ranked 581 of 14,804 drugs, putting it in the 95th percentile of treatments for asthma, and asthma was ranked 143 of 6,481 diseases, or in the 97th percentile of indications for imatinib. We note that other models can potentially improve the ranking performance compared to our simplistic logistic regression-based approach. However, those models lack mechanistic insight and ease of interpretability provided by our method.

Simply looking at the model’s top 10 paths that connect imatinib and asthma does not immediately reveal why imatinib could treat asthma. However, one path passes through the compound masitinib, and this compound is stated as treating asthma (Figure 7a). Masitinib is also a tyrosine kinase inhibitor that has been shown to have some effect on reducing asthma symptoms (Humbert *et al*., 2009). As both imatinib and masitinib are tyrosine kinase inhibitors, examining their common targets yields cKIT, a proto-oncogene that plays a role in acute myeloid leukemia (Gari *et al*., 1999; Edling and Hallberg, 2007). Filtering on cKIT, we find that in the top 10 paths, three paths that pass the concept mast cell leading to biological processes associated with asthma mast cell activation, histamine secretion by mast cell, and prostaglandin production involved in inflammatory response (Figure 7b). The role of the mast cell in the pathophysiology of asthma is well established and inhibiting activation of these cells could be one plausible mechanism for imatinib’s efficacy (Bradding *et al*., 2006; Amin, 2012). Reexamining the top 10 paths irrespective of target yields a path through “mast cell degranulation” where imatinib is purported to affect this process. Combining the results from these two sets of paths (Figures 7a & 7b) yields a more general and equally plausible mechanism, where cKIT inhibition by imatinib, prevents cKIT’s function in mast cell activation and degranulation, which in turn promotes asthma. This mechanism found through examining the top weighted paths in MechRepoNet is highly consistent with that previously (Cahill *et al*., 2017).

**Figure 7:**
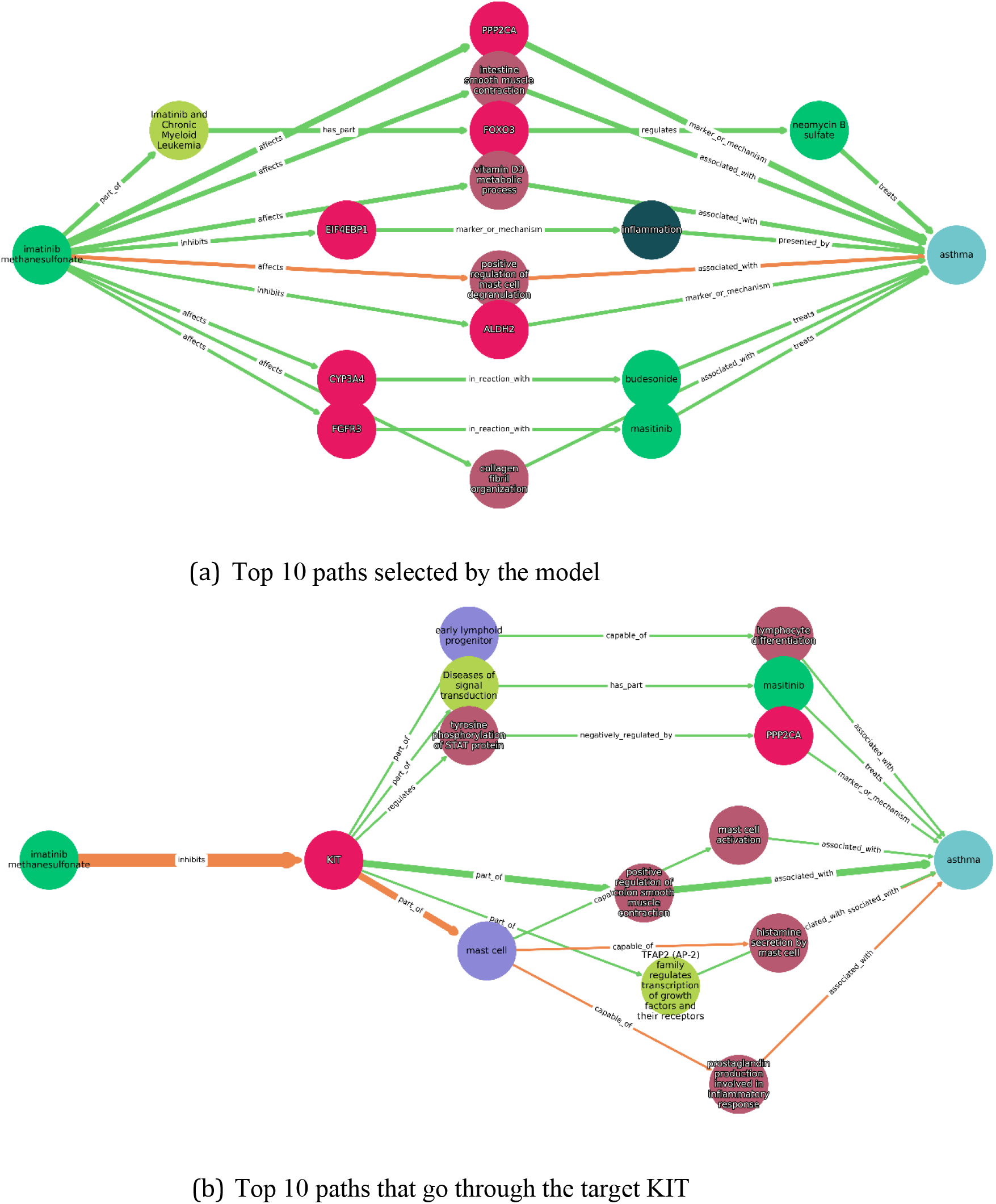
Top weighted paths from imatinib to asthma. The paths with the 10 largest weights connecting imatinib and asthma either (a) overall or (b) through cKIT. Paths mentioned in the text have been highlighted in orange.

## Discussion

Arguably more important than the computed confidence of a computational drug repurposing prediction is the reasoning behind that prediction. The reasoning chain provides a human interpretable explanation as to what mechanisms could be at play when producing a repurposing prediction. Guided by a mechanistic prediction, a domain expert would be better able to assess a prediction’s evidence than just the model’s probability score alone, and even guide further experimental validation of the hypothesis. If our goal were to produce the absolute best classifier for drug repositioning with maximized evaluation metrics, a model that includes the much more predictive similarity metapaths, or possibly a more abstract deep learning model would have been preferable (Zhu *et al*., 2020). However, this weaker predictor, being entirely based on paths with mechanistic meaning, provides a level of human interpretability not otherwise present.

Our method outlined in this work is not only able to rank potential repurposing candidates but also provide important biological context to the results. Each path identified through this method consists of relationships from multiple sources, joined together through common concepts. The data sources selected for integration individually have a high level of curation and therefore contain extensive knowledge regarding individual drugs and diseases. However, the large amount of data produces numerous multi-step paths from any single drug to a disease. Through well-engineered features derived from true drug mechanisms, our repurposing model can identify the patterns in this data most likely to be important in a treatment context.

We recognize that a limit to MechRepoNet’s mechanistic interpretability is its underlying data, as not all features from DrugMechDB are represented in our final model. Our analysis demonstrates that a significant challenge in computational repurposing based on knowledge graphs is the presence of significant gaps in the underlying knowledge graphs. Despite designing MechRepoNet based on the edges found in DrugMechDB, we still only found 20 out of 123 (16%) paths to be completely represented in MechRepoNet, demonstrating that many key mechanistic relationships are not readily available in structured databases. We believe this type of analysis is useful to guiding future curation efforts aimed at drug repurposing.

Our case study identified informative paths that potentially warrant follow-up experimentation and further research. In these paths, every link represents a testable hypothesis that can be verified through experimentation, or some new potential avenue of treatment that can be explored. These predictive models can aid in research by providing new hypotheses about a drug’s connections to a disease. Finally, our method is generalizable and could be applied to other applications such as to predict drug-phenotype or drug-physiological process pairs and its mechanistic rationalization for personalized medicine.

## Acknowledgements

This work was supported by the National Institutes of Health [R01 AG066750, OT2 TR003427].

